# Serotonin inhibits sexual receptivity in female cichlid fish

**DOI:** 10.1101/2025.01.09.632154

**Authors:** Paige Lee, Domino Joyce

## Abstract

Serotonin is involved in multiple diverse and complex biological processes. Selective serotonin reuptake inhibitors (SSRIs) are often used to test this, and fluoxetine, a commonly detected SSRI in aquatic ecosystems from pharmaceutical pollution, has been found to disrupt important behaviours and functions in organisms exposed to the antidepressant. Increased serotonin generally has an inhibitory effect on sexual behaviour and decreased serotonin has a facilitatory effect, however in the few existing studies on fish, either no effect or the opposite effect have been reported in females, with most studies focusing on male behaviour. The present research aims to examine the role of the serotonin pathway in female sexual receptivity in the cichlid fish *Nyassachromis* cf. *microcephalus*. The sexual dimorphism and lekking system of this species, where male-male competition is aggressive and female choice is important, make them an important and interesting evolutionary system. Serotonin levels were modulated using fluoxetine and after a three-week exposure to their respective treatments (control - 0µg/L, low ecologically relevant dose - 0.54µg/L, high dose - 5.4µg/L), female sexual receptivity to male courtship was measured in a simulated lek mating arena where multiple males were presented simultaneously. Sexual receptivity was inhibited in females exposed to a high dose of fluoxetine. This behaviour change due to a disruption of the serotonin pathway could affect sexual selection in the long term, and in the short term highlights the potential for pollution to affect aquatic life by directly altering mating behaviour.

**Highlights:**

- Disrupting the serotonin pathway reduces sexual receptivity in female cichlid fish
- The inhibitory effect of fluoxetine on female receptivity is dose-dependent
- The serotonin pathway could be important in the evolution of male mating strategies
- SSRI pollution could cause behaviour changes that amplify freshwater biodiversity loss

## 1. Introduction

Serotonin, also known as 5-hydroxytryptamine (5-HT), is a neurotransmitter that is involved in multiple biological processes across many species. It affects locomotion in molluscs [1,2] and nematodes [3,4], feeding in annelids [5,6] and crustaceans [7,8], learning and memory in flies [9,10] and fish [11,12], social behaviour in mice [13,14] and monkeys [15,16], sexual behaviour in birds [17,18] and rats [19,20], and more. Its roles are diverse and complex. Considering sexual behaviour alone, serotonin is involved in altering male attraction to females in Eurasian starlings *Sturnus vulgaris* [18], sexual receptivity in female fruit flies *Drosophila melanogaster* [21], sexual motivation and copulatory behaviours in male Wistar rats *Rattus norvegicus* [19,22], territorial aggression in male bower-building cichlids *Nyassachromis* cf. *microcephalus* [23], and courtship behaviour in male Mediterranean field crickets *Gryllus bimaculatus* [24].

A variety of drugs that impact the serotonergic system have been used to study the links between serotonin and sexual behaviour [25,26], most extensively selective serotonin reuptake inhibitors (SSRIs) which are a class of antidepressants. They block or reduce the reabsorption of 5-HT by the presynaptic neuron, retaining more 5-HT in the synapses and enhancing 5-HT transmission to postsynaptic receptors [27,28]. Evidence in general suggests that increased 5-HT inhibits sexual behaviour and decreased 5-HT facilitates sexual behaviour [29,30], with most studies focusing on male response [22,29,31], some studies exploring the combined effects on males and females [32–34] and more recently how females responded [30,35,36] and the differential effects on males and females [20,37,38].

The role of serotonin in female sexual behaviour is also important to understand in the context of environmental pollution, since antidepressants are widely prescribed and administered throughout the world, and enter the ecosystem through human or animal excretions and the disposal of unwanted drugs [39]. Fluoxetine in particular, has increasingly been detected in water bodies [40], and Brooks et al. (2005) [41] found the presence of fluoxetine, sertraline and their metabolites in fish tissues sampled from an effluent-dominated stream. While the level of fluoxetine found in the water systems do not exceed the hazardous threshold [42], these environmentally relevant concentrations still disrupt important traits and behaviours in fish [43–46].

Existing studies on the effect on female fish have mostly shown no changes in behaviour (e.g. fathead minnow *Pimephales promelas* [47] and guppies *Poecilia reticulata* [48]) which is surprising since serotonin does regulate reproductive function in female fish [49], affecting the release of the gonadotropin-release hormone (GnRH) and in turn, the secretion of follicle-stimulating hormone (FSH) and luteinizing hormone (LH), which are involved in the development of ovaries and egg production in females. Changes in behaviour, especially sexual receptivity, might therefore be expected alongside these functional modulations in female fish. Sexual receptivity in females enables or encourages reproductive success, i.e. egg fertilisation, through their willingness to accept mating attempts from males [50,51]. Variation already exists due to factors such as hormonal systems and sexual experience [52,53], and female receptivity in turn influences mate preference and mating strategies [54,55].

From an evolutionary point of view, female receptivity may increase or decrease female fitness depending on the mating system of the species in question and a better understanding of the underlying mechanisms for sexual receptivity will enable its role in sexual selection to be further explored. Understanding how the serotonin pathway fits into the dynamics of sexual receptivity could provide insights into how sexual selection may be impacted. While it seems that females are typically less receptive when serotonin levels are increased, its effect can be dependent on other factors such as age and male behaviour. The present research investigates the relationship between serotonin and female sexual receptivity to male courtship by using the SSRI fluoxetine to modulate serotonin levels in female lekking cichlids *Nyassachromis* cf. *microcephalus*.

The haplochromine cichlids of Lake Malawi make an excellent model system with which to study this, since they are sexually dimorphic and their evolutionary history has been shaped by sexual selection [56,57]. *Nyassachromis* cf. *microcephalus* is one of many species where males build sand-castle “bowers” in breeding ground leks to attract females and defend them against intruder males [58]. When females visit the lek, males approach them and perform courtship displays. Females may choose to follow a male to his bower and circle with him, before spawning eggs that she turns around to collect in her mouth together with sperm released by the male. Being mouthbrooders, females leave the lek holding these fertilised eggs in their buccal cavity and release their offspring as free-swimming juveniles approximately three weeks later. Individuals can abandon the mating behaviour sequence at any time [59]. The distinct courtship behaviours by males and pre-spawning behaviours by females make *N*. cf. *microcephalus* an ideal species to examine female sexual receptivity, and the lekking system of this cichlid where male-male competition is aggressive and female choice is fundamental provides an opportunity to examine mate preference as a variable [59,60]. Young et al. (2009) [59] observed that females responded more favourably to males with larger bowers and Stauffer et al. (2005) [61] discovered in another lekking species that males with larger bowers enjoyed greater success in terms of eggs laid by females. Furthermore, the neural pathways that underpin sexual motivation in teleost fishes—including the serotonin pathway—are thought to be evolutionarily conserved [62], making this species highly suitable for researching the role of serotonin in sexual receptivity. This study tested the prediction that fluoxetine changes female sexual receptivity at environmentally relevant and high concentrations, in a simulated lek mating arena with multiple bowers and males presented simultaneously.

## 2. Methods

A double blind behaviour trial with video recordings was conducted to test whether female receptivity to male courtship changed when females were exposed to fluoxetine.

### 2.1. Subjects

The study was conducted between December 2022 and January 2023 in the aquarium at the University of Hull, using 28 female *Nyassachromis* cf. *microcephalus* descended from wild-caught populations from Lake Malawi in Africa. Females of at least 70mm standard length (nose to peduncle) were randomly selected from the stock tanks to maximise likelihood of them being sexually mature and receptive during the experiment. Individuals were anaesthetised using tricaine methanesulfonate (MS-222) before Passive Integrated Transponder (PIT) tags were inserted into their abdominal cavity, and standard length and weight were measured.

Subjects were then assigned by a third person to one of three neighbouring treatment tanks in turn after tagging. These tanks were of dimensions 177.5cm(L) x 45cm(B) x 39cm(H) and monitored for 24 hours to confirm recovery of subjects.

Each tank had a polypropylene filter grid propped up in the centre by a brick, a plant pot and half a plant pot to familiarise females with swimming through a barrier, as well as an air driven sponge filter to maintain water quality (see Figure 1). The barriers measured 40cm x 34cm, were 1.5cm thick, and each square grid was either 1.8cm or 3.6cm wide.

**Figure 1.**
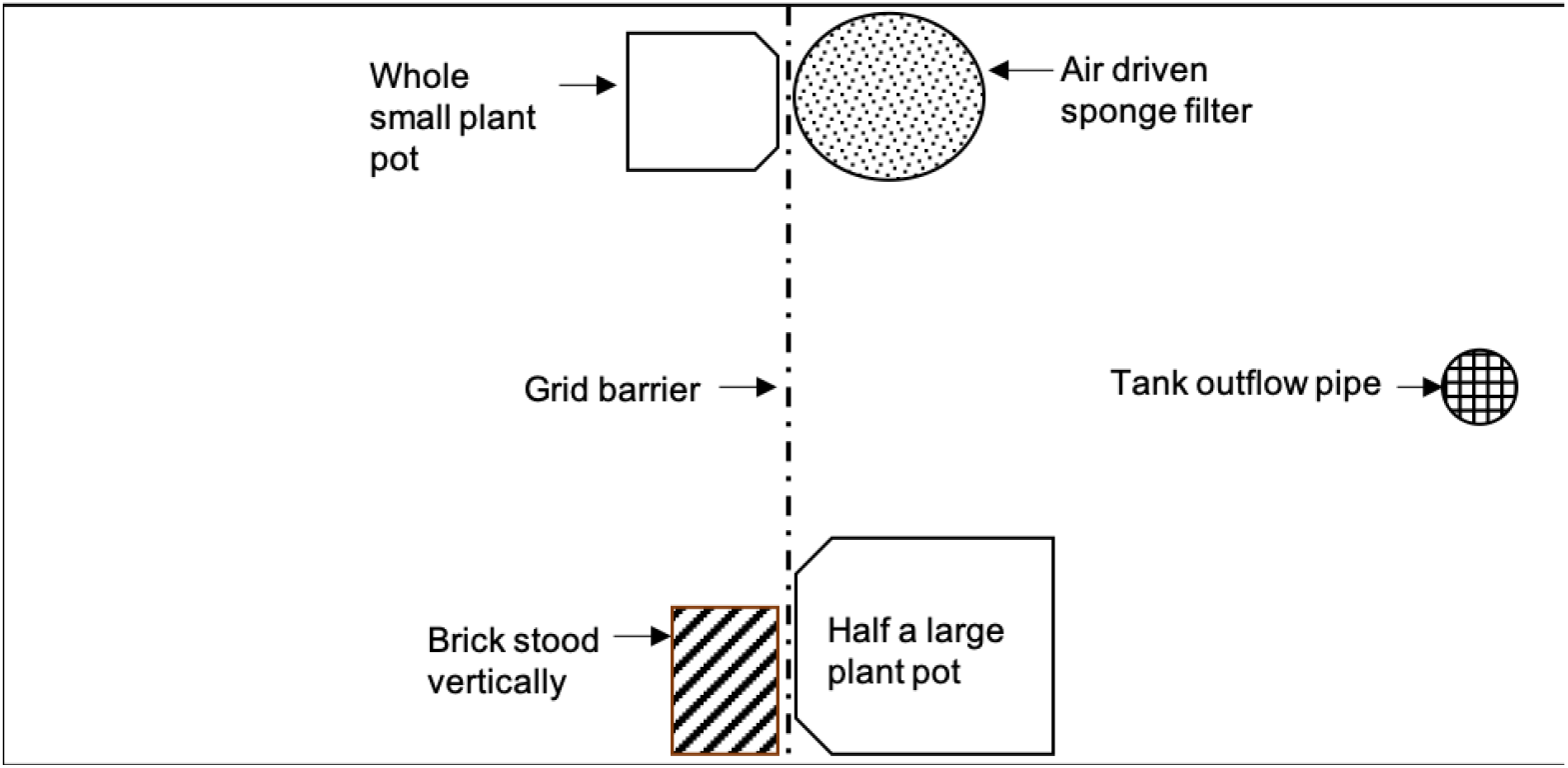
Schematic diagram of treatment tank set up viewed from above. Plant pots and brick served to hold the grid barrier in place and also as shelter or enrichment for the subjects.

During the treatment period when fluoxetine solution was administered, the treatment tanks were taken off the aquarium flow-through system and the water volume standardised at 215 litres. Water temperature was maintained at 24°C with an ambient temperature of 25°C and a 12:12 light-dark cycle in the aquarium. The cichlids received a varied diet of ZM (Zebramanagement) and Vitalis pellets once daily. They were fed approximately 0.5% of the total body mass in the tank.

### 2.2. SSRI treatment

The selective serotonin reuptake inhibitor (SSRI) used in this study was fluoxetine, purchased in hydrochloride form (#F375072, Fluorochem). Every seven days, a fresh stock solution of 540mg/L concentration was prepared by dissolving 54mg of fluoxetine hydrochloride in 100mL of purified water and stored in amber glass bottles at 4°C. This stock solution was then further diluted to produce the desired concentrations for each treatment group.

The three treatment tanks were each randomly assigned an SSRI treatment using a random sequence generator [63] and the fish in each tank were dosed on day 0 by a third person. For subjects assigned to the high dose treatment group (n=10), 2.15mL of the fluoxetine stock solution was added to their tank using a micropipette to produce a concentration of 5.4µg/L. For subjects assigned to the low (environmentally relevant) dose treatment group (n=11), 0.215mL of stock solution was added to their tank to produce a 0.54µg/L concentration. Subjects assigned to the control group (n=7) had 1.18mL of water (average of the amounts administered to the other two treatment groups) pipetted into their tank. These concentrations were selected following reports on fluoxetine presence in the environment [64,65] and literature on fluoxetine experiments with cichlids [43,66].

The cichlids were exposed to their respective treatment for 21 days to ensure that each individual could complete at least one reproductive cycle [67] and be more likely to be sexually receptive during the experimental period that followed. A refreshment dose was administered to the treatment tanks every 72 hours (Days 3, 6, 9, 12, 15, 18, 21). This schedule was derived from literature reporting the maximum absorption of fluoxetine to occur three days into the exposure period for Japanese medaka *Oryzias latipes* [68].

The amount of refreshment dose administered to each tank was determined by calculating the change in quantity of chemical in water over time using the equation introduced by Barron et al. (1990) [69]:

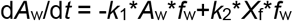

The variables in the equation were estimated following the methods from Lee et al. (2022) [23], but with fish weight *f*w being the total body mass in each treatment tank where individual fish weight was assumed to be constant throughout the experiment.

Prior to each refreshment dose, water tests for pH, ammonia and nitrite were conducted to ensure adequate water quality.

### 2.3. Experimental procedure

On day 21, the subjects were transferred from the three treatment tanks to an experimental arena tank containing untreated water. They were given 36 hours to acclimatise and explore their new environment before the experimental period started on day 23. The experimental tank measured 780cm(L) x 57cm(B) x 36cm(H) and was maintained on the aquarium flow-through system at a water level of 30.5cm. This tank was partitioned into seven segments using polypropylene filter grid barriers identical to the ones used in the treatment tanks (see Figure 2). These barriers were kept in place using bricks.

**Figure 2.**
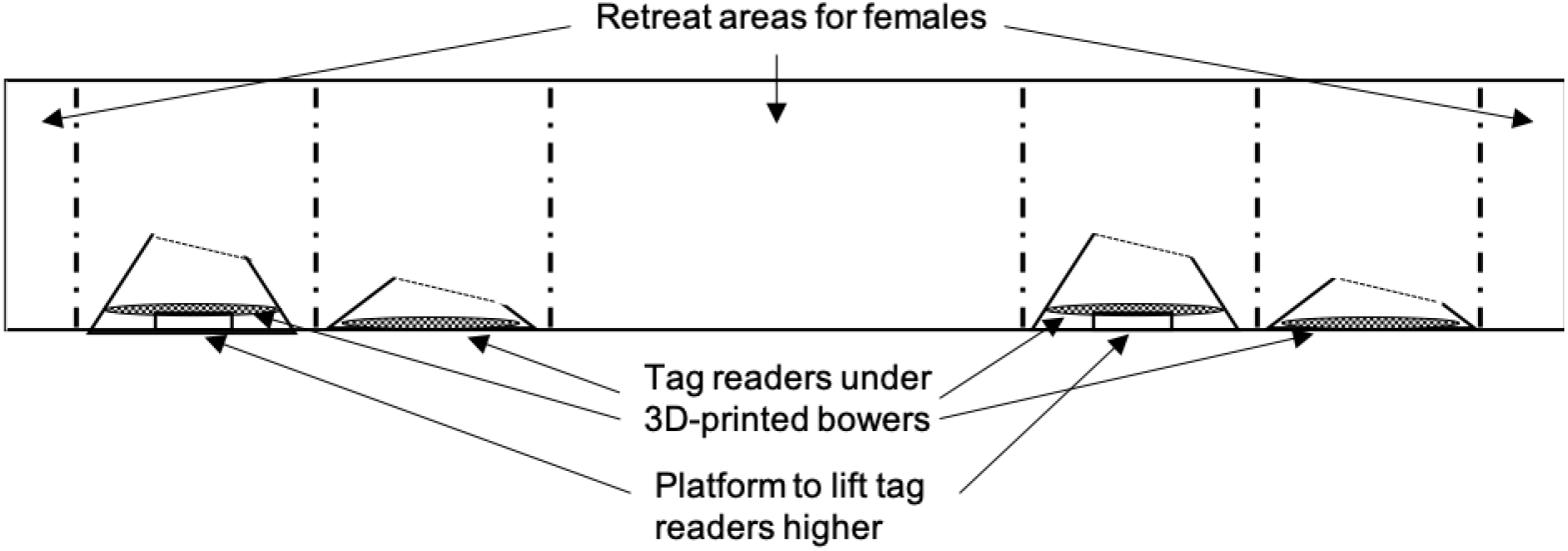
Schematic diagram of experimental tank set up.

Four tank segments of dimensions 78cm(L) x 57cm(B) x 36cm(H) were designated “bower segments”. Each bower segment contained an artificial bower 3D-printed using acrylonitrile butadiene styrene (ABS plastic) and spray painted a sandy colour using Plasti Dip Camo aerosol spray in Tan. These hollow artificial bowers were modelled after the shape of *N*. cf *microcephalus* sand bowers measured in Lake Malawi [70] and two bower sizes were created digitally: 1) a tall bower measuring 49.2cm at its widest diameter, 40.6cm at its narrowest diameter, and 12cm in height, and 2) a short bower identical to the control bower except scaled to a height of 6cm.

Two tall bowers and two short bowers were then printed, painted and randomly assigned to the four bower segments of the experimental tank (see supplementary Figure S1) using a random sequence generator [63]. On day 14, four sexually mature and dominant, i.e. vibrantly coloured, *N*. cf *microcephalus* males individually identified by PIT tags (previously used in [23]), and of a similar size (maximum of 10% difference in standard length) were selected from the stock tanks and randomly assigned a bower segment in the experimental tank using a random sequence generator [63]. They were given seven days to establish their territory in the bower segment before the focal females were transferred to the experimental tank on day 21. The larger grids (3.8cm wide) in the polypropylene barriers allowed females to swim through but not the larger males to swim through. The three remaining tank segments measuring 156cm(L) x 57cm(B) x 36cm(H) served as areas for females to retreat to if required.

Each bower segment contained a PIT tag reader placed under the bower. The tag readers allowed subjects from all treatments to be tested at the same time simulating a lek and avoiding testing order as a potential confounding factor. For the tall bowers, tag readers were placed on five plant pot saucers stacked together to form an approximately 6cm high platform such that the distance between the tag reader and the top of the bower were uniform across the tall and short bowers. These tag readers were connected by cables to a DEC-MPD-8 Multi-Point Decoder (MPD) run using SPDLogtags [71]. This software was run continuously throughout the experimental period to record the detection of individual fish that passed within 12cm of the tag reader and bower via their PIT tags (tag data). Tag data was exported as a csv file at the end of each day for conducting statistical analyses.

Four cameras were set up on tripods facing the experimental tank and connected to a 4-channel Swann DVR9-1425 CCTV recorder. These cameras recorded activity from 8:30am to 12:30pm during the experimental period on day 23 and 24. At the end of each day, video recordings were transferred onto an external hard drive and backed up on a cloud storage service (OneDrive) for carrying out behavioural observations.

### 2.4. Behavioural observations

Behaviour was scored using BORIS (Behavioral Observation Research Interactive Software) [72], following an ethogram derived from preliminary observations of *N*. cf *microcephalus* behaviour (Table 1). The behaviour of each male was observed and recorded (keys 0-2; see Table 1) from 8:30am to 12:30pm on days 23 and 24, i.e. within three days of the female subjects’ last exposure to their respective treatments on day 21. Where a female responded to the behaviour of a male during this period, tag data was used to identify the female, and her behaviour in response to the male would be observed and recorded (keys 3-9; see Table 1). Out of the 28 females in the experimental tank, 13 females were identified and observed. While this meant that half the total subjects were not observed, repeating the experiment would have involved either exposing the current subjects to their treatment again for 21 days (after ensuring there was no longer any fluoxetine in their system) without a guarantee that the remaining females would be identified and observed, or using a new group of subjects and increasing the number of fish used. In line with the “reduction” and “refinement” principles of the 3Rs of animal research [73], the study proceeded with a final number of 13 subjects included in the analysis.

**Table 1.** Ethogram used for behavioural observations of male and female cichlids.

| Key | Behaviour code | Description | Behaviour type |
| --- | --- | --- | --- |
| 0 | dance display | male swims in circles or figure of eight pattern around or in view of focal female while displaying upright dorsal fin and downwards pointing pectoral fin | state event |
| 1 | quiver display | male quivers body close to bower surface while displaying side to focal female with upright dorsal fin and downwards pointing pectoral fin | state event |
| 2 | chase | male accelerates towards focal female | point event |
| 3 | watch | female hovers in place within 10cm of male, facing him level in the water | point event |
| 4 | follow | female swims after male towards bower territory (above bower) | point event |
| 5 | visit | female follows male into bower territory (above bower) | point event |
| 6 | circle | female swims in a circle with male with bottom of body close to or brushing bower surface | state event |
| 7 | peck | female mouth open or close, making contact with bower | point event |
| 8 | avoid | female turns or swims away from male | point event |
| 9 | shoal | female hovers or swims with other females | state event |

The behaviour data collected was exported as a csv file for conducting statistical analyses, and it was only at this stage that the researchers learnt the treatment group of each subject. This allowed behavioural observations to be carried out without bias from knowing the experimental conditions beforehand.

### 2.5. Statistical analyses

Generalised linear mixed models (GLMMs) were fitted for the frequency of circling behaviour per female as a measure of sexual receptivity (key 6; see Table 1; circling is highly correlated with spawning [74]), testing for 1) treatment and bower height as fixed effects whilst controlling for male courtship frequency as a random effect to account for individual differences between males; and 2) treatment as a fixed effect while controlling for bower height and male courtship frequency as random effects. Male courtship frequency is maintained as a random effect as the main interest of this study is in the quantitative response of females to treatment. A comparison of Akaike Information Criterion (AIC) was used to select the most appropriate models (see supplementary Tables S1 and S2). All statistical analyses were conducted using R v4.2.2 [75] in RStudio v2024.9.1.394 [76], with the packages *tidyverse* v2.0.0 [77], *cowplot* v1.1.3 [78], *DT* v0.33 [79], *glmmTMB* v1.1.9 [80], *MuMIn* v1.48.4 [81], *performance* v0.12.4 [82], *DHARMa* v0.4.7 [83], *car* v3.1-3 [84], *multcomp* v1.4-26 [85], *ggsignif* v0.6.4 [86] and *ggpubr* v0.6.0 [87].

### 2.6. Reproducibility

Videos from the experimental period may be accessed by emailing the corresponding authors; behavioural observations were carried out using these videos on BORIS [72] and this project file is available at: https://doi.org/10.6084/m9.figshare.27635658. All data generated by and relating to this study are available at: https://doi.org/10.6084/m9.figshare.27635109. All code for the statistical analyses conducted on RStudio [76] and the resulting output file are available at: https://doi.org/10.6084/m9.figshare.27635643.

### 2.7. Ethical approval

Work was carried out with approval from the University of Hull AWERB and Faculty Ethics Committee, under UK Home Office Project license number PP0536451.

Sample size was kept at the minimum to be informative, in line with 3Rs guidance and ARRIVE guidelines (see supplementary material).

## 3. Results

Accounting for fluoxetine treatment and bower height as fixed effects and male courtship frequency as a random effect in a negative binomial GLMM, bower height was not found be a significant predictor of female circling frequency (x^2^=0.0096, df=1, p=0.922; see supplementary Figure S2). Bower height was therefore tested as a random effect together with male courtship frequency, but the best fit model accounted for only male courtship frequency as a random effect and treatment as a fixed effect. In a negative binomial GLMM, fluoxetine treatment was found to be a significant predictor of female circling behaviour (x^2^=6.62, df=2, p=0.0366).

Tukey’s post hoc test indicated a significantly lower frequency of circling behaviour in the high dose group compared to the control group (mean difference=1.99 occurrences per hour, p=0.0273; see Figure 4). Despite females in the low dose group also circling less than the females in the control group (see Figure 4), this difference was not significant (mean difference=1.51 occurrences per hour, p=0.287). Females in the low dose and high dose groups did not differ significantly in the frequency of circling behaviour displayed (mean difference=0.48 occurrences per hour, p= 0.559).

**Figure 3.**
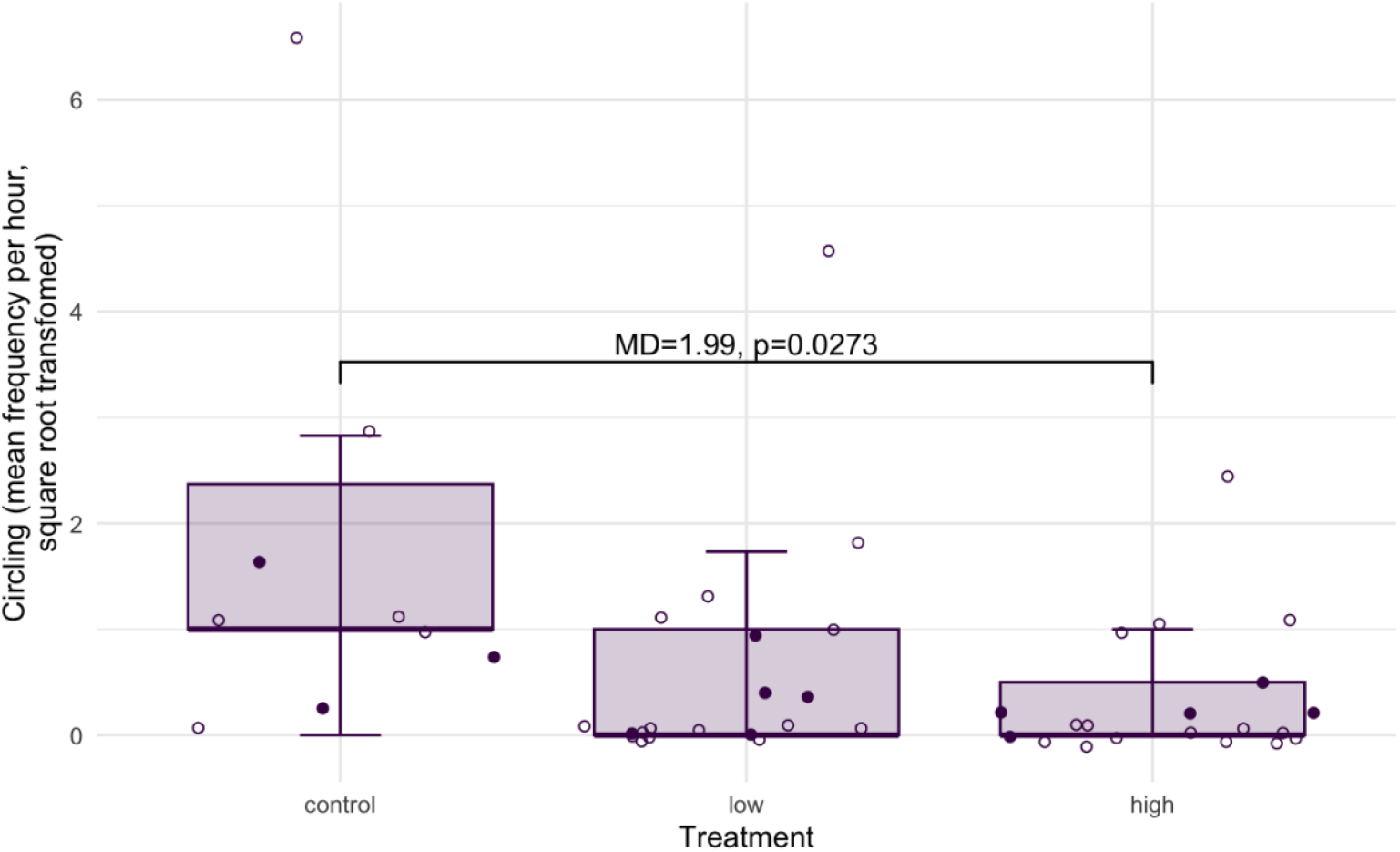
Female circling frequency differed significantly between treatments (x2=6.62, df=2, p=0.0366), with the high dose group displaying significantly less circling behaviour compared to the control group. The solid markers represent mean circling frequency per female, with individual observations at each bower as clear markers; see Table 2 for descriptive statistics.

**Table 2.** Summary of circling frequency per hour for each treatment group.

| Treatment | N | Median | Min | Max | Q1 | Q3 | IQR | Mean | SD | SE |
| --- | --- | --- | --- | --- | --- | --- | --- | --- | --- | --- |
| control | 3 | 1.12 | 0.12 | 5.38 | 0.62 | 3.25 | 2.63 | 2.21 | 2.79 | 1.61 |
| low | 5 | 0.38 | 0 | 2.62 | 0 | 0.5 | 0.5 | 0.7 | 1.1 | 0.49 |
| high | 5 | 0.12 | 0 | 0.75 | 0.12 | 0.12 | 0 | 0.22 | 0.3 | 0.13 |

A similar activity pattern was observed in the tag read count data collected, with the control group detected much more frequently at the bowers compared to the low and high dose treatment groups (see supplementary Figure S4). A positive correlation between behaviour data and tag data was confirmed (see supplementary Figure S5), which suggests that tag data may be used as an approximation of mating behaviour for future studies.

## 4. Discussion

The present study tested whether SSRI exposure alters female sexual receptivity in the lekking cichlid *Nyassachromis* cf. *microcephalus*, finding that compared to untreated females, fluoxetine-treated females indeed engaged in fewer circling events with males in response to courtship, though this difference was significant only for the high dose group (5.4µg/L fluoxetine) and not the low, environmentally relevant dose group (0.54µg/L fluoxetine). To the extent of the authors’ knowledge, this demonstrates for the first time in fish, the inhibitory effect of serotonin on female sexual receptivity.

Existing literature has established that increased serotonin levels (as would be expected with SSRI treatment) inhibit female sexual receptivity in various taxa [18,30,88,89], but this relationship has thus far not been reported in teleosts. Studies report either a lack of relationship [47,48] or the inverse relationship. In the Siamese fighting fish *Betta splendens*, females treated with 54μg/L fluoxetine (but not those treated with 0.54μg/L fluoxetine) for 7 days engaged in significantly longer spawning sessions compared to untreated females [36], and in the killifish *Nothobranchius furzeri*, females engaged in significantly more mating events per spawning session when treated with 5.3μg/L fluoxetine (but not those treated with 0.7μg/L fluoxetine) for 14 weeks compared to untreated females [90].

Whilst the treatment effect on *B. splendens* and *N. furzeri* is the opposite of that for *N*. cf. *microcephalus*, it identifies other variables that may influence the relationship between serotonin and female sexual receptivity. *Nothobranchius furzeri*, juveniles were dosed soon after hatching until adulthood whereas the *N*. cf. *microcephalus* were dosed as adults. In rodents, fluoxetine also had contrasting effects on Sprague-Dawley rats *Rattus norvegicus* depending on the life stage at which the drug was administered, where treatment of neonatal females facilitated receptivity once they were sexually mature [91] while treatment of adult females inhibited receptivity [92]. In the same studies, the respective authors found that the inhibition of receptivity in adult females was accompanied by disruptions to the reproductive system in the form of elongated estrous cycles but surprisingly, developmental fluoxetine exposure did not alter estrous cycles later in life despite the increase in receptive behaviours; the exact mechanisms triggered by increased serotonergic activity early in life that led to this reported effect in adult female rats are not fully understood yet.

This variation in how serotonin impacts reproduction is also evident in teleosts, as Thoré et al. [90] discovered in killifish that the increase in female sexual receptivity correlated with an increase in egg production while Forsatkar et al. [36] reported a decrease in egg production in Siamese fighting fish despite the increase in female sexual receptivity. In separate studies on fathead minnows *Pimephales promelas*, Weinberger and Klaper [47] found that serotonin did not affect female sexual receptivity and Parrott and Metcalfe [93] later found that serotonin increased egg production. In cichlids, no studies assessing the role of serotonin in both female reproductive axis and sexual receptivity have so far been reported, however Latifi et al. [43] revealed that fluoxetine exposure decreased egg production in the convict cichlid *Amatitlania nigrofasciata* while Dorelle et al. [94] found that fluoxetine exposure increased LH levels (associated with increased sexual receptivity) in the cichlid *Cichlasoma dimerus*. Considering the varied and often contrasting effects between species and within the reproductive axis, e.g. oocyte maturation [95,96], fertilisation rate [97,98], and estradiol levels [97,99], broadening the investigation on the current study species to examine any changes in reproductive functions that might occur alongside the inhibitory effect of serotonin on sexual receptivity identified here will contribute towards a fuller picture of the relationship between serotonin, sexual behaviour and reproduction in fish.

In *Betta splendens*, the concentration of fluoxetine subjects were exposed to was ten times higher than that for *N*. cf. *microcephalus* albeit for a shorter period, which suggest sensitivities to exposure duration and dosage. Acute and chronic exposures have been suggested to be typically 96 hours at most and at least 10% of a species’ lifespan respectively [100], however these durations are not standardised across experimental procedures [101]. Chronic exposure to fluoxetine can range from 21 days [102] to months [103], and while the 7-day exposure period for *B. splendens* was not categorised as either, the duration falls closer to acute exposure. The 21-day exposure period for *N*. cf. *microcephalus* could be considered chronic but a longer exposure period following the recommendation in Newman (2014) [100] would be ideal for gaining a better understanding of the long-term effects of fluoxetine on female receptivity, especially at environmentally relevant concentrations for which there was no significant effect observed in the present study. Fergusson et al. [103] exposed *P. reticulata* to fluoxetine for a period representing 2-3 overlapping generations, with the mean generation time estimated to be around 7 months [104] and 30% of their approximate lifespan [105]. While the lifespan of the *N*. cf. *microcephalus* in the wild has not yet been reported, the generation time for *Nyassachromis microcephalus* is estimated to be 1-2 years [106] (although observations from the lab indicate that the *N*. cf. *microcephalus* in captivity start to breed in less than a year) and future investigations into the chronic effects of SSRI pollution on *N*. cf. *microcephalus* behaviour could use this as a guide for determining exposure duration.

In addition to varying across species, exposure timing in relation to life stage and exposure duration, the influence of SSRIs on female sexual behaviour also varies with dosage. The dose-dependent effects of fluoxetine reported in this study highlight a sex difference in sensitivity to changes in serotonin levels within *N*. cf. *microcephalus*. In females, sexual receptivity was only inhibited when exposed to 5.4µg/L fluoxetine while in males, fluoxetine exposure at a concentration of 0.54µg/L was sufficient to significantly reduce territorial aggression [23], warranting further dose-effect experiments to better understand the impact potential changes in environmental SSRI levels—which are very likely considering global trends in mental health issues and SSRI consumption [107,108]—could have on each sex and how these differential responses might interact.

In a lekking species where the mating strategy involves male-male competition and female choice, reduced aggression in dominant bower-building males with SSRI exposure [23] would reduce their effectiveness in defending their territories, giving subordinate males increased chances of sneaking copulations with females without the costs of building and maintaining bowers [67]. As bower size has been posited to have a role in female mate preference [59,109], the current study had aimed to also determine how SSRI exposure would affect that. However, the best fit statistical model for this dataset discounted bower height as a factor when comparing female mating frequency between experimental treatments, suggesting that male courtship is more important than bower height in female mating decisions. Therefore, if the inhibition of female receptivity due to SSRI exposure means that courtship by dominant males becomes less relevant in achieving mating success, this again could give subordinate males increased chances of reproducing as the role of female choice in this mating system is diluted, and alternative reproductive tactics could emerge.

Overall, this study illustrates the importance of understanding the serotonin pathway’s role in both female sexual receptivity and ultimately, perhaps the evolution of mating strategies. From an applied perspective, this study demonstrates the potential for pollution to affect aquatic life by directly altering mating behaviour, which could exacerbate biodiversity loss in freshwater environments.

## CRediT authorship contribution statement

**PL:** Conceptualization, Data curation, Formal analysis, Investigation, Methodology, Project administration, Resources, Visualization, Writing – original draft, Writing – review and editing. **DJ:** Conceptualization, Funding acquisition, Methodology, Project administration, Resources, Supervision, Validation, Writing – review and editing.

## Declaration of interests

The authors declare that they have no known competing financial interests or personal relationships that could have appeared to influence the work reported in this paper.

## Acknowledgements

We would like to thank Stuart Butterick, Georgia Melbourne, Nathaniel Brown, and Louise France, for their assistance in refining and printing the 3D bower models, Alan Smith for fish husbandry advice, and AS and Owen Walsh for technical support. We are grateful to the EvoHull Lab members for further logistical support and discussions throughout the study, James Gilbert for statistical advice, and Lesley Morrell for additional supervision.

## Funding

This work was supported by the University of Hull as part of the Happy Chemical PhD scholarship cluster awarded to DJ.

## Notes

### Competing Interest Statement

The authors have declared no competing interest.

