## Supplementary Material for "Serotonin inhibits sexual receptivity in female cichlid fish"

#### Supplementary Methods

Artificial bower creation:
First, an artificial physical bower model was crafted using concrete and wire mesh. Next, photos were taken of this bower and used in photogrammetry software Regard3D [1] to create a digital model. The digital model was further refined using Autodesk Meshmixer [2]. Two bower shapes were printed and used in this study (Figure S1).

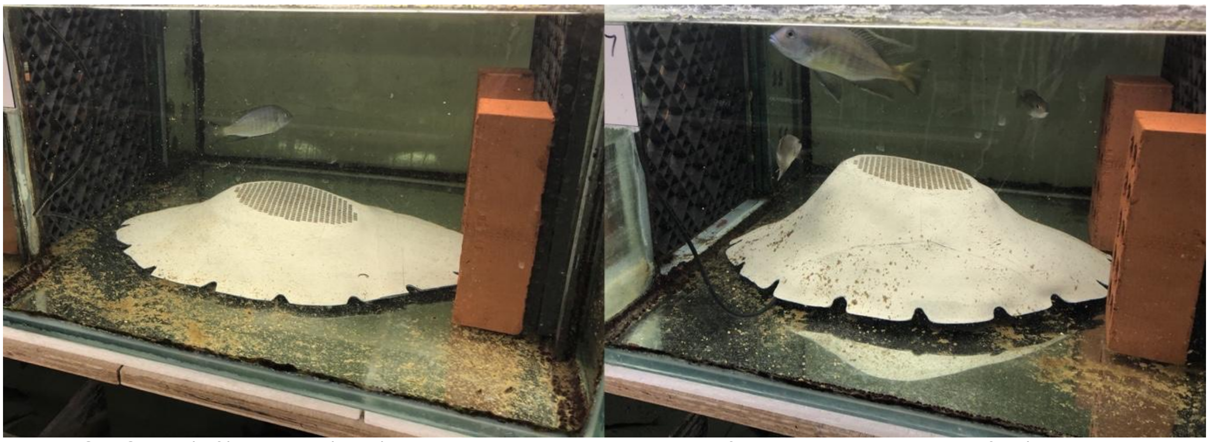

*Figure S1. Short (left) and tall (right) 3D-printed bowers with the platform converted to a grid for (ultimately unsuccessful) egg collection and gaps around the bottom edges to equalise water pressure inside and outside the bower model.*

Fitting statistical models:
Poisson GLMMs were initially fitted for female circling frequency, testing for treatment and bower height as fixed effects whilst controlling for male courtship frequency as a random effect. The best fit model (delta AICc = 5.64) excluded the interaction of the fixed effects (see Table S1).

*Table S1. Poisson model comparison table for female circling frequency with treatment and bower height as fixed effects and male courtship frequency as a random effect, ranked by AICc value.*

| Fixed effects interaction | Random effects | df | logLik | AICc | delta | weight |
|---|---|---|---|---|---|---|
| no | yes | 5 | -95.82 | 203.6 | 0 | 0.944 |
| yes | yes | 7 | -95.64 | 209.3 | 5.64 | 0.056 |
| no | no | 4 | -113.38 | 236.1 | 32.43 | 0 |
| yes | no | 6 | -112.53 | 240.0 | 36.33 | 0 |

Poisson GLMMs were then fitted to test for only treatment as a fixed effect whilst controlling for bower height as a random effect together with male courtship frequency. The best fit model (delta AICc = 2.71) included only male courtship frequency as a random effect (see Table S2).

*Table S2. Model comparison table for female circling frequency with treatment as a fixed effect, ranked by AICc value.*

| Random effects | df | logLik | AICc | delta | weight |
|---|---|---|---|---|---|
| male courtship frequency | 4 | -95.82 | 200.9 | 0 | 0.795 |
| male courtship frequency + bower height | 5 | -95.82 | 203.6 | 2.71 | 0.205 |
| bower height | 4 | -117.91 | 245.1 | 44.19 | 0 |

Due to the presence of overdispersion and zero-inflation in these Poisson models, negative binomial models were ultimately fitted for the final analyses. Simulated residuals confirmed satisfactory fits.

#### Supplementary Results

Circling frequency and tag reader detections at bowers:
Female circling frequency did not differ significantly between short and tall bowers (see Figure S2).

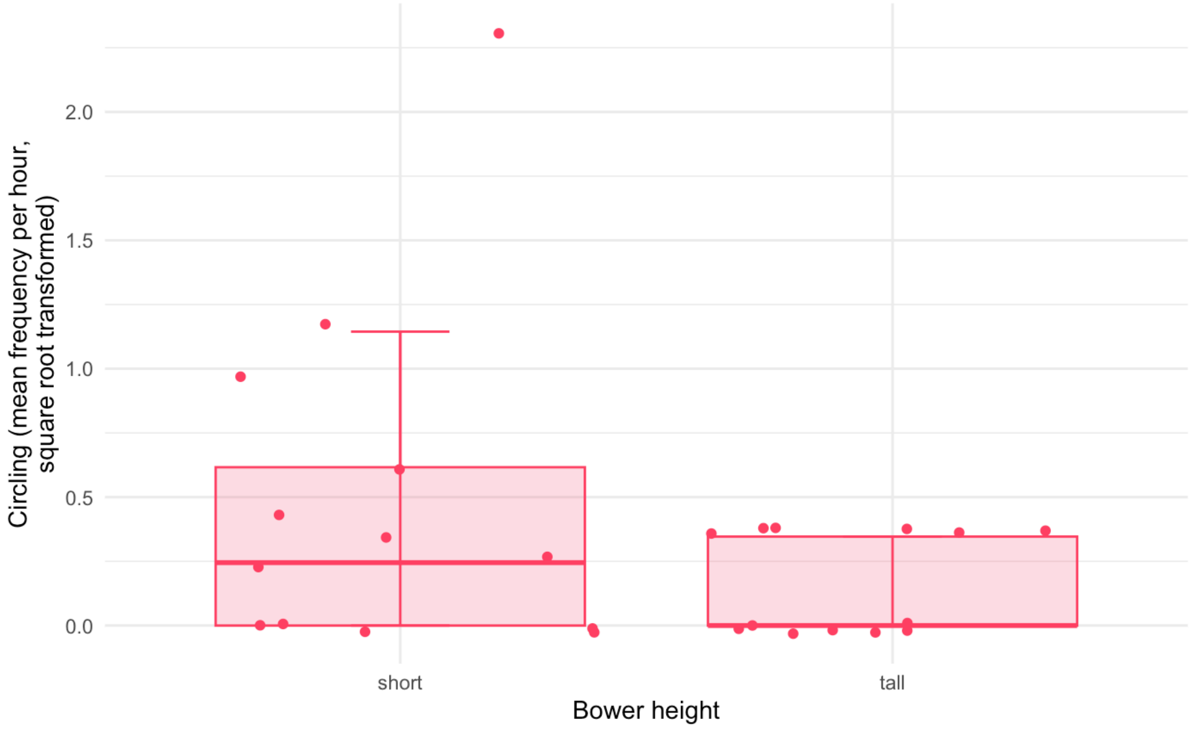


*Figure S2. Female circling frequency did not differ significantly between tall and short bowers ($x^2$=0.0096, df=1, p=0.922). The round markers represent the frequency of circling displays for each subject.*

The tag data collected during the experimental period was examined. While females in the low dose treatment group reflected a similar pattern of females being detected more frequently at short bowers compared to tall bowers, females in the control group and high dose group appeared to be more active at tall bowers compared to short bowers (see Figure S3).

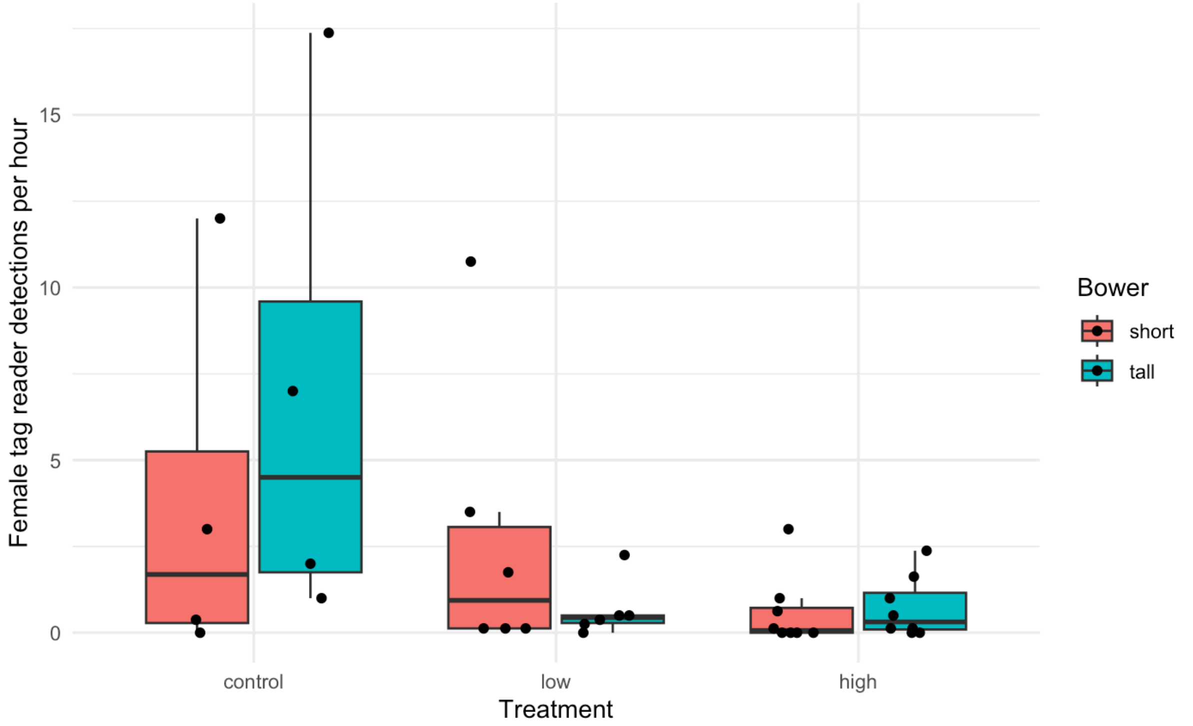


*Figure S3. Tag data shows more female activity detected at tall bowers compared to short bowers for the control group and high dose group, although the difference is less pronounced for the high dose group and there was generally more activity detected for the control group. In contrast, more female activity was detected at short bowers compared to tall bowers for the low dose group. The round markers represent the number of tag reader detections for each subject.*

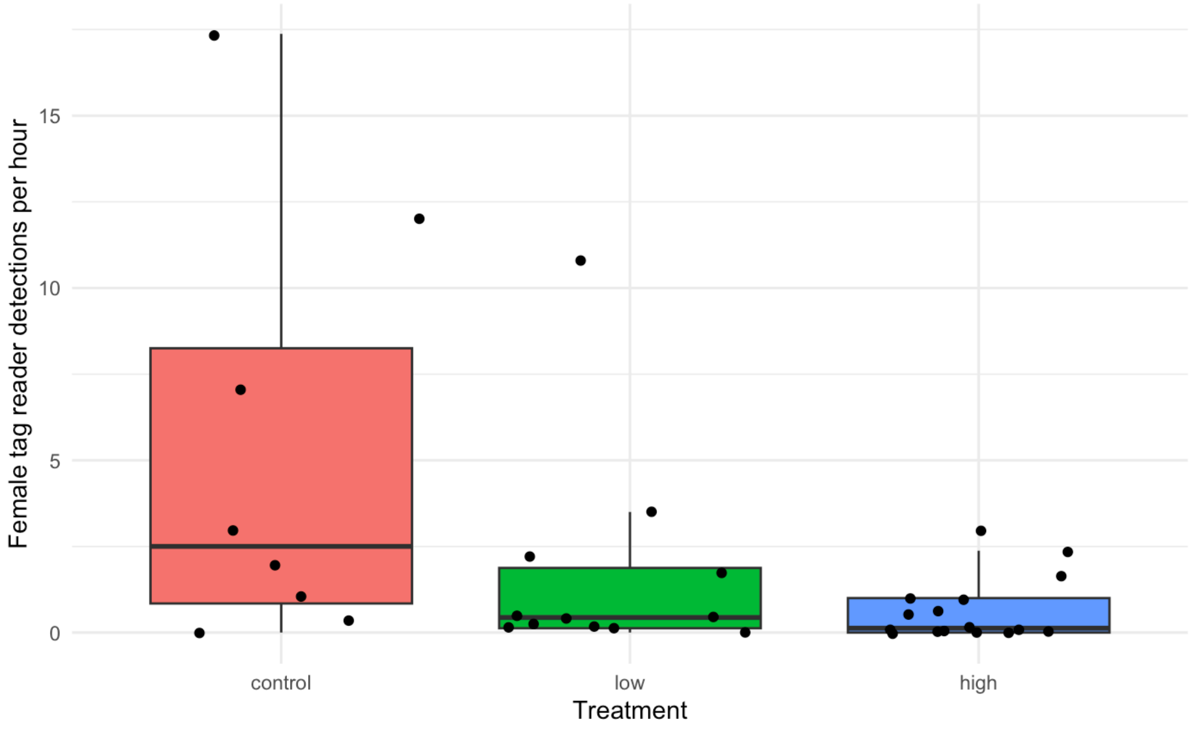

*Figure S4. Tag data shows females in the control group were detected most frequently, followed by females in the low dose group and females in the high dose group were detected least frequently. The round markers represent the number of tag reader detections for each subject.*

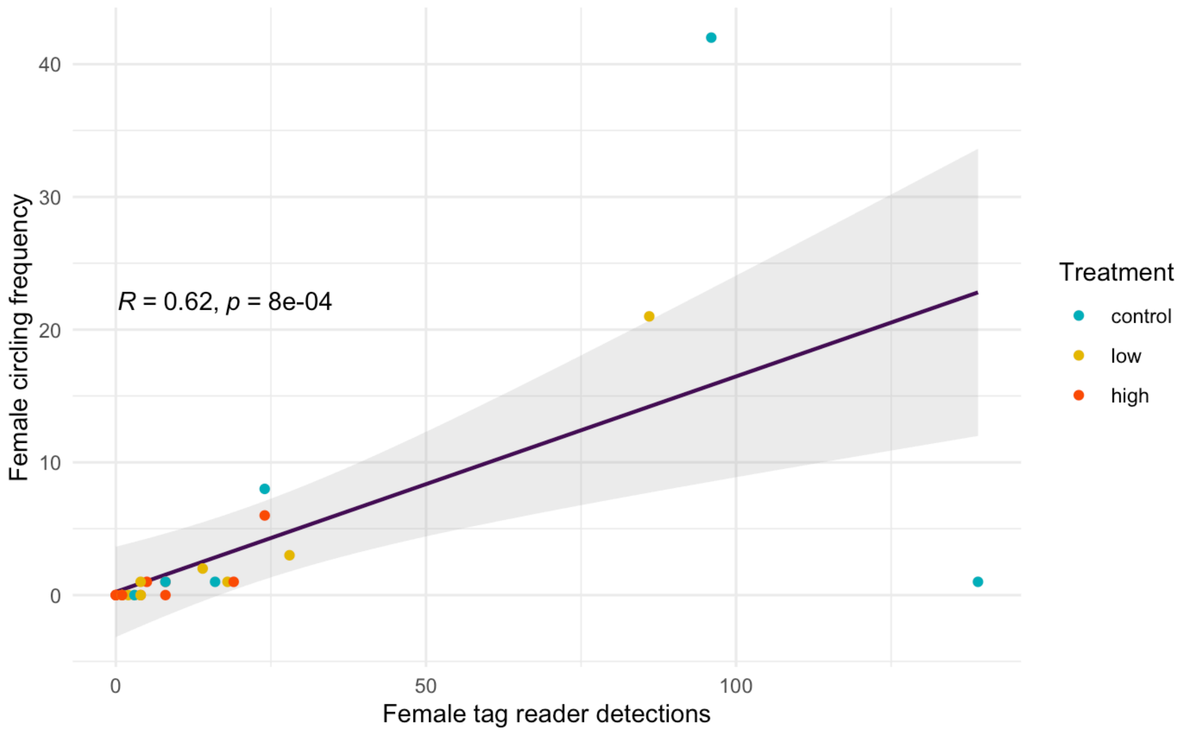


*Figure S5. The frequency of tag detections correlates positively with the frequency of circling behaviour by the subjects at tall and short bowers (r=0.62, df=18, p<0.001), which are represented by the round markers.*

NOTE: Please save this file locally before filling in the table, DO NOT work on the file within your internet browser as changes will not be saved. Adobe Acrobat Reader (available free here) is recommended for completion.

### The ARRIVE guidelines 2.0: author checklist

#### The ARRIVE Essential 10

These items are the basic minimum to include in a manuscript. Without this information, readers and reviewers cannot assess the reliability of the findings.

| Item | | Recommendation | Section/line number, or reason for not reporting |
|---|---|---|---|
| **Study design** | 1 | For each experiment, provide brief details of study design including: | |
| | | a. The groups being compared, including control groups. If no control group has been used, the rationale should be stated. | Section 2.1, 2.2 |
| | | b. The experimental unit (e.g. a single animal, litter, or cage of animals). | Section 2.1 |
| **Sample size** | 2 | a. Specify the exact number of experimental units allocated to each group, and the total number in each experiment. Also indicate the total number of animals used. | Section 2.1, 2.2, 2.4 |
| | | b. Explain how the sample size was decided. Provide details of any *a priori* sample size calculation, if done. | Section 2.1, 2.4 |
| **Inclusion and exclusion criteria** | 3 | a. Describe any criteria used for including and excluding animals (or experimental units) during the experiment, and data points during the analysis. Specify if these criteria were established *a priori*. If no criteria were set, state this explicitly. | Section 2.1, 2.4 |
| | | b. For each experimental group, report any animals, experimental units or data points not included in the analysis and explain why. If there were no exclusions, state so. | Section 2.4 |
| | | c. For each analysis, report the exact value of *n* in each experimental group. | Section 2.2, and Line 350 |
| **Randomisation** | 4 | a. State whether randomisation was used to allocate experimental units to control and treatment groups. If done, provide the method used to generate the randomisation sequence. | Section 2.1, 2.2 |
| | | b. Describe the strategy used to minimise potential confounders such as the order of treatments and measurements, or animal/cage location. If confounders were not controlled, state this explicitly. | Section 2.3 |
| **Blinding** | 5 | Describe who was aware of the group allocation at the different stages of the experiment (during the allocation, the conduct of the experiment, the outcome assessment, and the data analysis). | Section 2.1, 2.2, 2.3, 2.4 |
| **Outcome measures** | 6 | a. Clearly define all outcome measures assessed (e.g. cell death, molecular markers, or behavioural changes). | Section 2.3, 2.4 |
| | | b. For hypothesis-testing studies, specify the primary outcome measure, i.e. the outcome measure that was used to determine the sample size. | Section 2.4 |
| **Statistical methods** | 7 | a. Provide details of the statistical methods used for each analysis, including software used. | Section 2.5 |
| | | b. Describe any methods used to assess whether the data met the assumptions of the statistical approach, and what was done if the assumptions were not met. | Section 2.5, 2.6, 3 |
| **Experimental animals** | 8 | a. Provide species-appropriate details of the animals used, including species, strain and substrain, sex, age or developmental stage, and, if relevant, weight. | Section 2.1, 2.3 |
| | | b. Provide further relevant information on the provenance of animals, health/immune status, genetic modification status, genotype, and any previous procedures. | Section 2.1, 2.3 |
| **Experimental procedures** | 9 | For each experimental group, including controls, describe the procedures in enough detail to allow others to replicate them, including: | Section 2 |
| | | a. What was done, how it was done and what was used. | Section 2 |
| | | b. When and how often. | Section 2.1, 2.2, 2.3 |
| | | c. Where (including detail of any acclimatisation periods). | |
| | | d. Why (provide rationale for procedures). | Section 2 |
| **Results** | 10 | For each experiment conducted, including independent replications, report: | |
| | | a. Summary/descriptive statistics for each experimental group, with a measure of variability where applicable (e.g. mean and SD, or median and range). | Section 3 |
| | | b. If applicable, the effect size with a confidence interval. | Section 3 |

### The Recommended Set

These items complement the Essential 10 and add important context to the study. Reporting the items in both sets represents best practice.

| Item | | Recommendation | Section/line number, or reason for not reporting |
|---|---|---|---|
| **Abstract** | 11 | Provide an accurate summary of the research objectives, animal species, strain and sex, key methods, principal findings, and study conclusions. | Line 25-44 |
| **Background** | 12 | a. Include sufficient scientific background to understand the rationale and context for the study, and explain the experimental approach. | Line 51-108 |
| | | b. Explain how the animal species and model used address the scientific objectives and, where appropriate, the relevance to human biology. | Line 110-134 |
| **Objectives** | 13 | Clearly describe the research question, research objectives and, where appropriate, specific hypotheses being tested. | Line 105-108, 131-134, 294-301 |
| **Ethical statement** | 14 | Provide the name of the ethical review committee or equivalent that has approved the use of animals in this study, and any relevant licence or protocol numbers (if applicable). If ethical approval was not sought or granted, provide a justification. | Section 2.7 |
| **Housing and husbandry** | 15 | Provide details of housing and husbandry conditions, including any environmental enrichment. | Section 2.1 |
| **Animal care and monitoring** | 16 | a. Describe any interventions or steps taken in the experimental protocols to reduce pain, suffering and distress. | Section 2.1, 2.2 |
| | | b. Report any expected or unexpected adverse events. | NA |
| | | c. Describe the humane endpoints established for the study, the signs that were monitored and the frequency of monitoring. If the study did not have humane endpoints, state this. | Section 2.1, 2.2, and Outlined in Home Office PPL No. PP0536451 |
| **Interpretation/ scientific implications** | 17 | a. Interpret the results, taking into account the study objectives and hypotheses, current theory and other relevant studies in the literature. | Section 4 |
| | | b. Comment on the study limitations including potential sources of bias, limitations of the animal model, and imprecision associated with the results. | Line 410-414, 423-429, 455-459 |
| **Generalisability/ translation** | 18 | Comment on whether, and how, the findings of this study are likely to generalise to other species or experimental conditions, including any relevance to human biology (where appropriate). | Line 431-437, 445-449, 460-465, 469-471 |
| **Protocol registration** | 19 | Provide a statement indicating whether a protocol (including the research question, key design features, and analysis plan) was prepared before the study, and if and where this protocol was registered. | Outlined in Home Office PPL No. PP0536451 |
| **Data access** | 20 | Provide a statement describing if and where study data are available. | Section 2.6 |
| **Declaration of interests** | 21 | a. Declare any potential conflicts of interest, including financial and non-financial. If none exist, this should be stated. | Line 482-484 |
| | | b. List all funding sources (including grant identifier) and the role of the funder(s) in the design, analysis and reporting of the study. | Line 477-478, 495-496 |
